# Detecting the impact of temperature on transmission of Zika, dengue and chikungunya using mechanistic models

**DOI:** 10.1101/063735

**Authors:** Erin A. Mordecai, Jeremy M. Cohen, Michelle V. Evans, Prithvi Gudapati, Leah R. Johnson, Catherine A. Lippi, Kerri Miazgowicz, Courtney C. Murdock, Jason R. Rohr, Sadie J. Ryan, Van Savage, Marta S. Shocket, Anna Stewart Ibarra, Matthew B. Thomas, Daniel P. Weikel

## Abstract

Recent epidemics of Zika, dengue, and chikungunya have heightened the need to understand the seasonal and geographic range of transmission by *Aedes aegypti* and *Ae. albopictus* mosquitoes. We use mechanistic transmission models to derive predictions for how the probability and magnitude of transmission for Zika, chikungunya, and dengue change with mean temperature, and we show that these predictions are well matched by human case data. Across all three viruses, models and human case data both show that transmission occurs between 18-34°C with maximal transmission occurring in a range from 26-29°C. Controlling for population size and two socioeconomic factors, temperature-dependent transmission based on our mechanistic model is an important predictor of human transmission occurrence and incidence. Risk maps indicate that tropical and subtropical regions are suitable for extended seasonal or year-round transmission, but transmission in temperate areas is limited to at most three months per year even if vectors are present. Such brief transmission windows limit the likelihood of major epidemics following disease introduction in temperate zones.

**Author Summary:** Understanding the drivers of recent Zika, dengue, and chikungunya epidemics is a major public health priority. Temperature may play an important role because it affects mosquito transmission, affecting mosquito development, survival, reproduction, and biting rates as well as the rate at which they acquire and transmit viruses. Here, we measure the impact of temperature on transmission by two of the most common mosquito vector species for these viruses, *Aedes aegypti* and *Ae. albopictus*. We integrate data from several laboratory experiments into a mathematical model of temperature-dependent transmission, and find that transmission peaks at 26-29°C and can occur between 18-34°C. Statistically comparing model predictions with recent observed human cases of dengue, chikungunya, and Zika across the Americas suggests an important role for temperature, and supports model predictions. Using the model, we predict that most of the tropics and subtropics are suitable for transmission in many or all months of the year, but that temperate areas like most of the United States are only suitable for transmission for a few months during the summer (even if the mosquito vector is present).

## Main Text

Epidemics of dengue, chikungunya, and Zika are sweeping through the Americas, and are part of a global public health crisis that places an estimated 3.9 billion people in 120 countries at risk [1]. Dengue virus (DENV) distribution and intensity in the Americas has increased over the last three decades, infecting an estimated 390 million people (96 million clinical) per year [2]. Chikungunya virus (CHIKV) emerged in the Americas in 2013, causing 1.8 million suspected cases from 44 countries and territories (www.paho.org). In the last two years, Zika virus (ZIKV) has spread throughout the Americas, causing 714,636 suspected and confirmed cases, with many more unreported (http://ais.paho.org/phip/viz/ed_zika_cases.asp, as of January 5, 2017). The growing burden of these diseases (including links between Zika infection and both microcephaly and Guillain-Barré syndrome [3]) and potential for spread into new areas creates an urgent need for predictive models that can inform risk assessment and guide interventions such as mosquito control, community outreach, and education.

Predicting transmission of DENV, CHIKV, and ZIKV requires understanding the ecology of the vector species. For these viruses the main vector is *Aedes aegypti*, a mosquito that prefers and is closely affiliated with humans, while *Ae. albopictus*, a peri-urban mosquito, is an important secondary vector [4,5]. We expect one of the main drivers of the vector ecology to be the climate, particularly temperature. For that reason, mathematical and geostatistical models that incorporate climate information have been valuable for predicting and responding to *Aedes* spp. spread and DENV, CHIKV, and ZIKV outbreaks [5–10].

The effects of temperature in ectotherms are largely predictable from fundamental metabolic and ecological processes. Survival, feeding, development, and reproductive rates predictably respond to temperature across a variety of ectotherms, including mosquitoes [11,12]. Because these traits help to determine transmission rates, the effects of temperature on transmission should also be broadly predictable from mechanistic models that incorporate temperature-dependent traits. Here, we introduce a model based on this framework that overcomes several major gaps that currently limit our understanding of climate suitability for transmission. Specifically, we develop models of temperature-dependent transmission for *Ae. aegypti* and *Ae. albopictus* that are (a) mechanistic, facilitating extrapolation beyond the current disease distribution, (b) parameterized with biologically accurate unimodal thermal responses for all mosquito and virus traits that drive transmission, and (c) validated against human dengue, chikungunya, and Zika case data across the Americas.

We synthesize available data to characterize the temperature-dependent traits of the mosquitoes and viruses that determine transmission intensity. With these thermal responses, we develop mechanistic temperature-dependent virus transmission models for *Ae. aegypti* and *Ae. albopictus*. We then ask whether the predicted effect of temperature on transmission is consistent with patterns of actual human cases over space and time. To do this, we validate the models with DENV, CHIKV, and ZIKV human incidence data at the country scale from the Americas from 2014-2016. To isolate temperature dependence, we also statistically controlled for population size and two socioeconomic factors that may influence transmission. If temperature fundamentally limits transmission potential, transmission should only occur at actual environmental temperatures that are predicted to be suitable, and conversely, areas with low predicted suitability should have low or zero transmission (i.e., false negative rates should be low). By contrast, low transmission may occur even when temperature suitability is high because other factors like vector control can limit transmission (i.e., the false positive rate should be higher than the false negative rate). Finally, if the simple mechanistic model accurately predicts climate suitability for transmission, then we can use it to map climate-based transmission risk of DENV, CHIKV, ZIKV, and other emerging pathogens transmitted by *Ae. aegypti* and *Ae. albopictus* seasonally and geographically.

## Results

### Temperature-dependent transmission

Data gathered from the literature [9,13–15,15–21,21–30] revealed that all mosquito traits relevant to transmission—biting rate, egg-to-adult survival and development rate, adult lifespan, and fecundity—respond strongly to temperature and peak between 23°C and 34°C for the two mosquito species (*Ae. aegypti* in Fig. 1 and *Ae. albopictus* in Fig. S1). DENV extrinsic incubation and vector competence peak at 35°C [31–37] and 31-32°C [31,32,34,38], respectively, in both mosquitoes—temperatures at which mosquito survival is low, limiting transmission potential (Figs. 1, S1). Appropriate thermal response data were not available for CHIKV and ZIKV extrinsic incubation and vector competence.

**Fig. 1.**
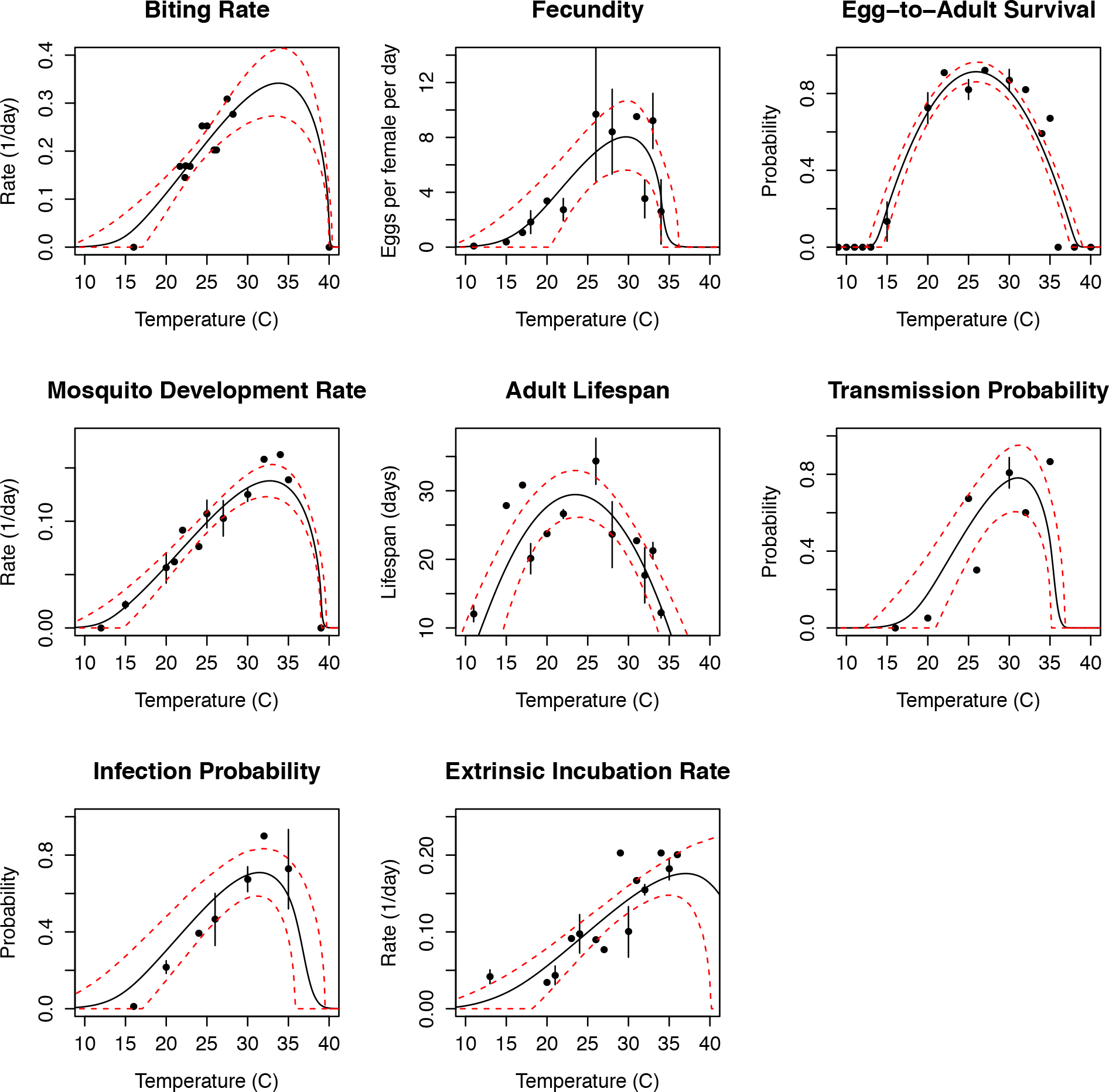
Thermal responses of *Ae. aegypti* and DENV traits that drive transmission (data sources listed in Table S2). Informative priors based on data from additional *Aedes* spp. and flavivirus studies helped to constrain uncertainty in the model fits (see Materials and Methods; Table S3). Points and error bars indicate the data means and standard errors (for display only; models were fit from the raw data). Black solid lines are the mean model fits; red dashed lines are the 95% credible intervals. Thermal responses for *Ae. albopictus* are shown in Fig. S1.

We estimated the posterior distribution of *R_0_*(*T*) and used it to calculate key temperature values that indicate suitability for transmission: the mean and 95% credible intervals (95% CI) on the critical thermal minimum, maximum, and optimum temperature for transmission by the two mosquito species. At constant temperature, *Ae. aegypti* transmission peaked at 29.1°C (95% CI: 28.4 – 29.8°C), and declined to zero below 17.8°C (95% CI: 14.6 – 21.2°C) and above 34.6°C (95% CI: 34.1 – 35.6°C) (Fig. 2). *Ae. albopictus* transmission peaked at 26.4°C (95% CI: 25.2 – 27.4°C) and declined to zero below 16.2°C (95% CI: 13.2 – 19.9°C) and above 31.6°C (95% CI: 29.4 – 33.7°C) (Fig. 2). Overall, the thermal response curve for *Ae. albopictus* is shifted towards lower temperatures than *Ae. aegypti*, so *Ae. albopictus* transmission is better suited to colder environments. For a more realistic scenario in which daily temperature ranged over 8°C, the transmission peak, minimum, and maximum were slightly lower for both *Ae. aegypti* (28.5°C, 13.5°C, 34.2°C, respectively) and *Ae. albopictus* (26.1°C, 11.9°C, and 28.3°C, respectively). The lower thermal maximum under fluctuating temperatures occurs because we incorporated empirically supported irreversible lethal effects of temperatures that exceed thermal maxima for survival (see Materials and Methods).

**Fig. 2.**
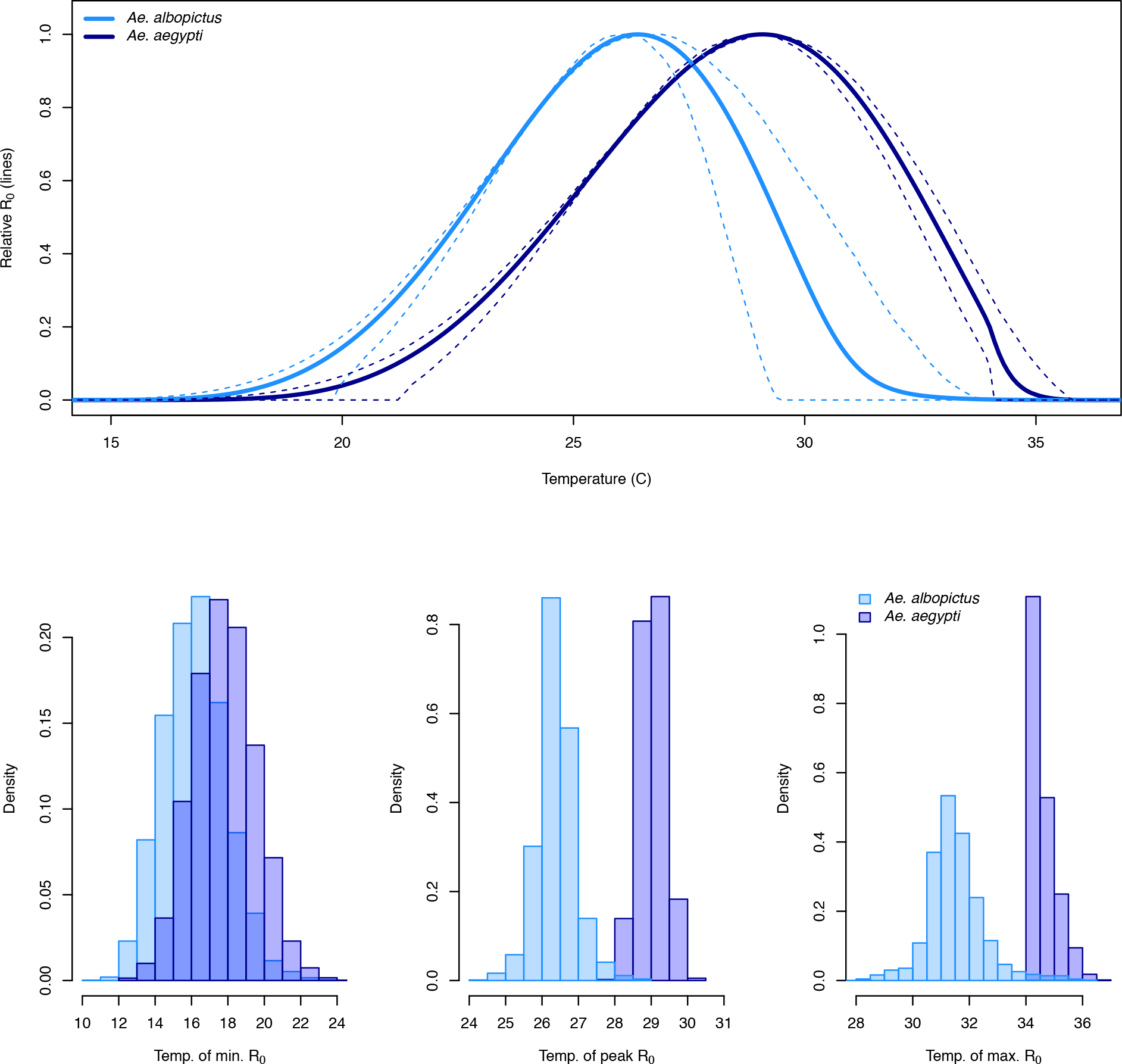
Relative *R_0_* across constant temperatures (°C; top) for *Ae. albopictus* (light blue) and *Ae. aegypti* (dark blue), and histograms of the posterior distributions of the critical thermal minimum (bottom left), temperature at peak transmission (bottom middle), and critical thermal maximum (bottom right; all in °C). Solid lines: mean posterior estimates; dashed lines: 95% credible intervals. *R_0_* curves normalized to a 0-1 scale for ease of comparison and visualization.

The posterior distribution of *R_0_*(*T*) allows us to evaluate uncertainty in key temperature values that define the transmission range, including critical thermal minimum, maximum, and optimum. Uncertainty was higher for the critical thermal minimum for transmission than for the maximum or optimum, and the two mosquito species overlapped most for this outcome (Fig. 2, bottom panels). This occurred because several trait thermal responses increase gradually from low to mid temperatures but decline more steeply at high temperatures (Fig. 1), so uncertainty is greatest at low temperatures. *Ae. aegypti* has a substantially higher optimum and maximum temperature than *Ae. albopictus* (Fig. 2) due to its greater rates of adult survival at high temperatures (see Supplementary Materials for sensitivity analyses).

### Model validation

We used generalized linear models (GLM) to ask whether the predicted relationship between temperature and transmission, *R_0_*(*T*), was consistent with observed human cases of DENV, CHIKV, and ZIKV. Specifically, we assessed whether *R_0_*(*T*) was an important predictor of the probability of autochthonous transmission occurring and of the incidence given that transmission occurred. We also controlled for human population size, virus species, and two socioeconomic factors. (Note that we focused on testing the *R_0_*(*T*) model, rather than on constructing the best possible statistical model of human case data.) To do this, we used the version of the *Ae. aegypti R_0_*(*T*) model that includes 8°C daily temperature range, along with country-scale weekly case reports of DENV, CHIKV, and ZIKV in the Americas and the Caribbean between 2014-2016. We first addressed the fact that countries with larger populations have greater opportunities for (large) epidemics by creating two predictors that incorporate scaled *R_0_*(*T*) and population size. In the models of the probability of autochthonous transmission occurring we used the product of the posterior probability that *R_0_*(*T*) > 0 (which we notate as *GR_0_*) and the log of population size (*p*) to give *log*(*p*)*\*GR_0_*. In the models of incidence given that transmission does occur we used the log of the product of the posterior mean of *R_0_*(*T*) and population size, *log*(*p*R_0_*(*T*)). To control for several socioeconomic factors that might obscure the impact of temperature, we also included log of gross domestic product (GDP) and log percent of GDP in tourism (using logs to improve normality). These are potential indicators of investment in and/or success of vector control and infrastructure improvements that prevent transmission. By comparing models that included the *R_0_*(*T*) metric alone, socioeconomic factors alone, or both, we tested whether *R_0_*(*T*) was an important predictor of observed transmission occurrence and incidence (see Table S4). Note that *R_0_*(*T*) is out of sample because it is derived and calculated strictly from laboratory data on mosquitoes, and we perform a validation analyses for *R_0_*(*T*) using independent case incidence reports. For this validation step we assessed model adequacy for the transmission data in two ways. First we used the full dataset for case incidence reports to select the best model (Table S4) and determine whether or not our predicted value of relative *R_0_*(*T*) based on laboratory data was included in the model (“within sample” analysis). Second we used a bootstrapping approach where models were fit on subsets of the case incidence data that were randomly sampled and then predictive accuracy of the competing models (Table S4) was assessed on left-out data (“out of sample” analysis).

For the probability of autochthonous transmission occurring, the model that included both the *R_0_*(*T*) predictor and socioeconomic predictors had overwhelming support based on Bayesian Information Criterion (BIC; model PA5 relative probability = 1, Table S4). Based on deviance explained, the models that included *R_0_*(*T*), with or without the socioeconomic predictors out-performed the model that did not include *R_0_*(*T*) (Table S4; Figs. 3A, S2). In analyses of out-of-sample accuracy, models that included the *R_0_*(*T*) metric (with or without the socioeconomic factors) were surprisingly accurate. They predicted the probability of transmission with 86-91% out-of-sample accuracy for DENV (Table S4). For CHIKV and ZIKV, models that included the *R_0_*(*T*) metric or population alone had 66-69% out-of-sample accuracy (Table S4). There were no significant differences in out-of-sample accuracy between the top four models but for both DENV and CHIKV/ZIKV the best model was significantly better than the worst model (see Supplementary Code for full results). The lower out-of-sample accuracy for CHIKV and ZIKV likely reflects the much lower frequency of positive values and the lower total sample size of this dataset. All results were similar for a set of models that separated *GR_0_* from population size, so for simplicity we show the model predictors that combines *GR_0_* and population size here (see Table S4 and Supplementary Code for results of other models). Further, from a biological perspective, the combined model better describes what we know about disease systems: if either the probability of *R_0_*(*T*) being greater than zero is small or population size is very small, transmission is unlikely to occur. Together, these analyses suggest that *R_0_*(*T*) is an important predictor of transmission occurrence, but that CHIKV and ZIKV need further data to better explain the probability of transmission occurrence (Figs. 3A, S2).

**Fig. 3.**
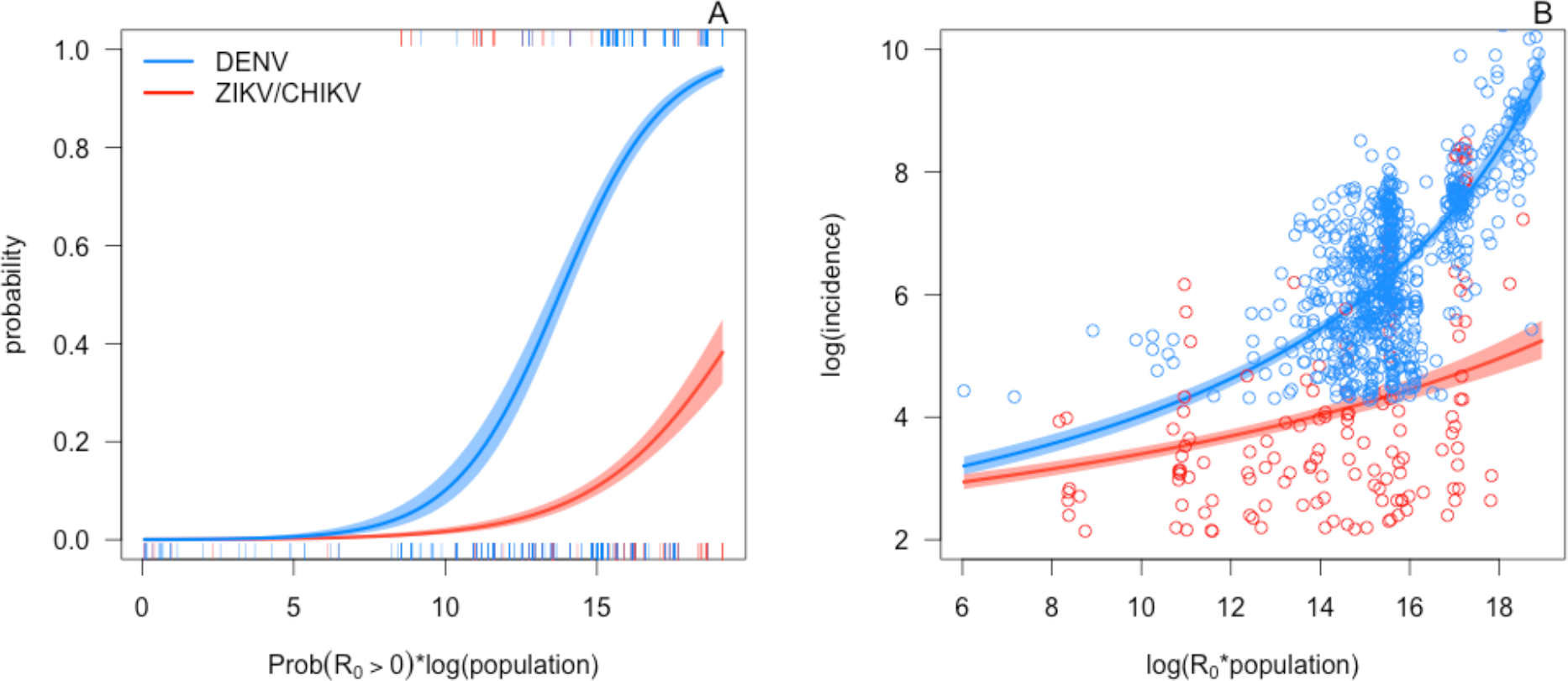
*Ae. aegypti R_0_*(*T*) and population size predict the probability and magnitude of transmission of DENV, CHIKV, and ZIKV across the Americas. A, *log*(*p*)*\*GR_0_* (the posterior probability that *R_0_*(*T*) > 0 times the log of population size) versus the probability of local transmission in the data. B, *log*(*p*R_0_*(*T*)) (log of *R_0_*(*T*) times the population size) versus the log of incidence, given that it exceeds the threshold for local transmission. Tick-marks and points: human transmission occurrence and incidence data, respectively, by country-week in the Americas and Caribbean. Lines and shaded areas: mean and 95% CI from GLM fits for DENV (blue) and CHIKV and ZIKV (red). For simplicity, we show the models that only include the covariates *log*(*p*)*\*GR_0_* or *log*(*p*R_0_*(*T*)), respectively, and do not include the socioeconomic covariates (models PA6 and IM4 in Table S4). For each case report data point, *log*(*p*)*\*GR_0_* and *log*(*p*R_0_*(*T*)) were calculated at the mean temperature 10 weeks prior to the reporting week [39].

*R_0_*(*T*) was also an important predictor of incidence, given that autochthonous transmission did occur. Within-sample, incidence was best predicted by the model that included both *R_0_*(*T*) and the socioeconomic predictors (model IM5 in Table S4) based on BIC (relative probability = 1). The models that included *R_0_*(*T*) out-performed those that did not based on deviance explained (Table S4). In out-of-sample validation, the models that included *R_0_*(*T*) explained the magnitude of incidence based on mean absolute percentage error (85-86% accuracy versus 83% accuracy for models that did not include *R_0_*(*T*); Table S4), but this difference was not statistically significant. For illustration, we show the simpler model that only contains the *R_0_*(*T*) predictor in the main text (Fig. 3B; model IM1 in Table S4). Notably, the models that contained *R_0_*(*T*) predicted incidence well for all three viruses, despite the lower incidence of CHIKV and ZIKV.

Although predicted *R_0_*(*T*) correlated with the observed occurrence and magnitude of human incidence for all three viruses, these observed incidence metrics were higher for DENV than for CHIKV and ZIKV. While the reason for this difference is unclear, the most likely explanation is that DENV is much more established in the region, so it is more likely to be detected, diagnosed, and reported. Because ZIKV and CHIKV are newly emerging, they may not have fully saturated the region at this early stage.

The ability of the model to explain the probability and magnitude of transmission is notable given the coarse scale of the human incidence versus mean temperature data (i.e., country-scale means), the lack of CHIKV- and ZIKV-specific trait thermal response data to inform the model, the nonlinear relationship between transmission and incidence, and all the well-documented factors other than temperature that influence transmission. Together, these analyses show simple mechanistic models parameterized with laboratory data on mosquitoes and dengue virus are consistent with observed temperature suitability for transmission. Moreover, the similar responses of human incidence of ZIKV, CHIKV, and DENV to temperature suggest that the thermal ecology of their shared mosquito vectors is a key determinant of outbreak location, timing, and intensity.

### Mapping climate suitability for transmission

The validated model can be used to predict where transmission is not excluded (posterior probability that *R_0_*(*T*) > 0, a conservative estimate of transmission risk). Considering the number of months per year at which mean temperatures do not prevent transmission, large areas of tropical and subtropical regions, including Puerto Rico and parts of Florida and Texas, are currently suitable year-round or seasonally (Fig. 4). These regions are fundamentally at risk for DENV, CHIKV, ZIKV, and other *Aedes* arbovirus transmission during a substantial part of the year (Fig. 4). Indeed, DENV, CHIKV, and/or ZIKV local transmission has occurred in Texas, Florida, Hawaii, and Puerto Rico (www.cdc.gov). On the other hand, many temperate regions experience temperatures suitable for transmission three months or less per year (Fig. 4), and the virus incubation periods in humans and mosquitoes restrict the transmission window even further. Temperature thus limits the potential for the viruses to generate extensive epidemics in temperate areas even where the vectors are present. Moreover, many temperate regions with seasonally suitable temperatures currently lack *Ae. aegypti* and *Ae. albopictus* mosquitoes, making vector transmission impossible (Fig. 4, black line). The posterior distribution of *R_0_*(*T*) also allows us to map months of risk with different degrees of uncertainty (e.g., 97.5%, 50%, and 2.5% posterior probability that that *R_0_* > 0), ranging from the most to least conservative (Fig. S4).

**Fig. 4.**
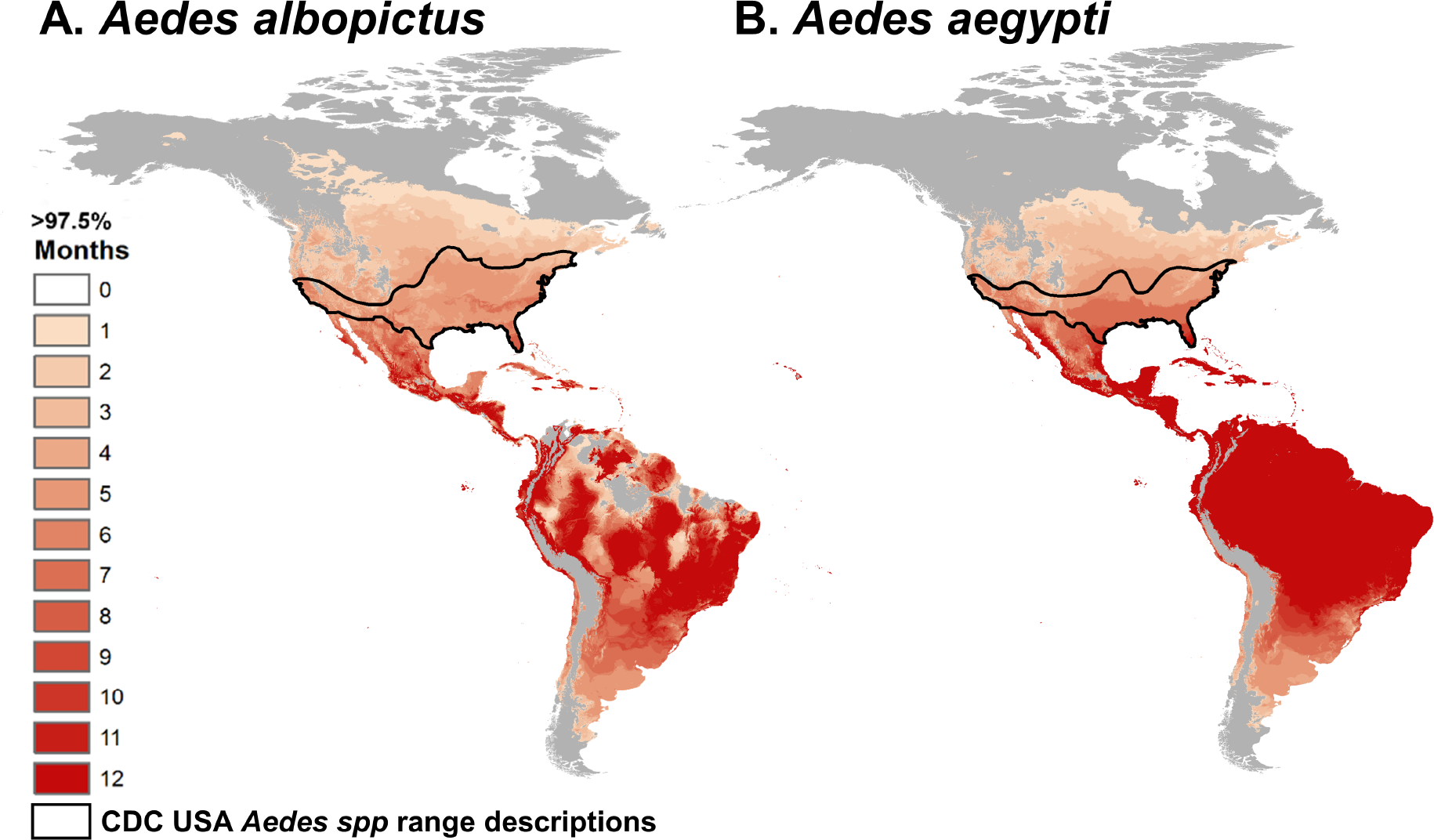
Map of predicted temperature suitability for virus transmission by *Ae. albopictus* and *Ae. aegypti*. Color indicates the consecutive months in which temperature is permissive for transmission (predicted *R_0_* > 0) for *Aedes* spp. transmission based on the minimum likely range (> 97.5% posterior probability that *R_0_* > 0). Black lines indicate the CDC estimated range for the two *Aedes* spp. in the United States. Model suitability predictions combine temperature mean and 8°C daily variation and are informed by laboratory data (Figs. 1, S1) and validated against field data (Fig. 3).

## Discussion

Temperature is an important driver of—and limitation on—vector transmission, so accurately describing the temperature range and optimum for transmission of DENV, CHIKV, and ZIKV is critical for predicting their geographic and seasonal patterns of spread [12,40]. We directly estimated the temperature – transmission relationship using mechanistic transmission models for each mosquito species (Fig. 2). These models are built using empirical estimates of the (unimodal) effects of temperature on mosquito and pathogen traits that drive transmission, including survival, development, reproduction, and biting rates (Figs. 1, S1). Because these trait thermal responses are unimodal across the majority of ectotherm taxa and traits, and because the traits combine nonlinearly to drive transmission, the emergent relationship between temperature and transmission is difficult to infer directly from field data or from individual trait responses. Here, we present a model of temperature-dependent DENV, CHIKV, and ZIKV transmission that advances on previous models because it is mechanistic, fitted from experimental trait data, and validated against independent human case data at a broad geographic scale (Fig. 3).

Mechanistic understanding is valuable for extrapolating beyond the current spatial and temporal range of transmission (Fig. 4), as compared to environmental niche models, for example [5,41,42]. Of the six previous mechanistic temperature-dependent models of DENV, CHIKV, or ZIKV transmission by *Ae. aegypti* and *Ae. albopictus* that we were able to reproduce, three had similar thermal optima [7,43,44] while the other three had dramatically higher optima (3-6°C) [9,45] (Fig. S5). Two models predicted much greater suitability for transmission at low temperatures [45], four predicted greater suitability at high temperatures [7,9,45], and two were very similar to ours [43,44] (Fig. S5). Only one of these previous models was (like ours) statistically validated against independent data not used to estimate model parameters, and its predictions were very similar to those of our model [43]. Other mechanistic and environmental niche models could not be directly compared with ours [5,10,40–42], either because fully reproducible equations, parameters, and/or code were not provided or because their predicted marginal effects of temperature were not displayed. Visually, our maps are similar to maps based on a previous model of *Ae. aegypti* and *Ae. albopictus* persistence suitability indices [40]. Recent environmental niche models of Zika distribution have shown similar but more constrained predicted distributions of environmental suitability, in part because these models include not just temperature suitability but also further environmental, socioeconomic, and demographic constraints [5,41,42,46].

Even though the thermal response data are imperfect—for example, CHIKV and ZIKV thermal response data are missing—and the human case data are reported at a coarse spatial scale, the validation analyses suggest that *R_0_*(*T*) is an important predictor of both the probability of transmission occurring and the magnitude of incidence for DENV, CHIKV, and ZIKV. This has several key implications. First, temperature-dependent transmission is pervasive enough to be detected at a coarse spatial scale. Second, dynamics of the mosquito predict transmission for a suite of *Ae. aegypti*-transmitted viruses, without additional virus-specific information. Third, climate and socio-economic factors combine to shape variation in incidence across countries. Finally, these simple predictors explain a substantial proportion of the variance in both the probability and intensity of transmission.

Predicting arbovirus transmission at a higher spatial resolution and precision will require more detailed information on factors like the exposure and susceptibility of human populations, environmental variation (e.g., oviposition habitat availability, seasonal and daily temperature variation), and socioeconomic factors. However, as a first step our mechanistic model provides valuable insight because it makes broad predictions about suitable environmental conditions for transmission, it is mechanistic and grounded in experimental trait data, it is validated against independent human case data, and its predictions are applicable across three different viruses. Using these thermal response models as a scaffold, additional drivers could be incorporated to obtain more precise and specific predictions about transmission dynamics, which could in turn be used for public health and vector control applications. For this purpose, all code and data used in the models are available as Supplementary Files.

The socio-ecological conditions that enabled CHIKV, ZIKV, and DENV to become the three most important emerging vector-borne diseases in the Americas make the emergence of additional *Aedes*-transmitted viruses likely (potentially including Mayaro, Rift Valley fever, yellow fever, Uganda S, or Ross River viruses). Efforts to extrapolate and to map temperature suitability (Fig. 4) will be critical for improving management of both ongoing and future emerging epidemics. Mechanistic models like the one presented here are useful for extrapolating the potential geographic range of transmission beyond the current envelope of environmental conditions in which transmission occurs (e.g., under climate change and for newly invading pathogens). Accurately estimating temperature-driven transmission risk in both highly suitable and marginal regions is critical for predicting and responding to future outbreaks of these and other *Aedes*-transmitted viruses.

## Materials and Methods

### Temperature-sensitive *R_0_* models

We constructed temperature-dependent models of transmission using a previously developed *R_0_* framework. We modeled transmission rate as the basic reproduction rate, *R_0_*—the number of secondary infections that would originate from a single infected individual introduced to a fully susceptible population. In previous work on malaria, we adapted a commonly used expression for *R_0_* for vector transmission to include the temperature-sensitive traits that drive mosquito population density:

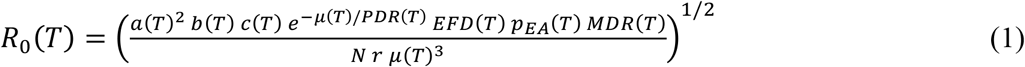

Here, (*T*) indicates that the trait is a function of temperature, *T*; *a* is the per-mosquito biting rate, *b* is the proportion of infectious bites that infect susceptible humans, *c* is the proportion of bites on infected humans that infect previously uninfected mosquitoes (i.e., *b*c* = vector competence), *µ* is the adult mosquito mortality rate (lifespan, *lf = 1/µ*), *PDR* is the parasite development rate (i.e., the inverse of the extrinsic incubation period, the time required between a mosquito biting an infected host and becoming infectious), *EFD* is the number of eggs produced per female mosquito per day, *p_EA_* is the mosquito egg-to-adult survival probability, *MDR* is the mosquito immature development rate (i.e., the inverse of the egg-to-adult development time), *N* is the density of humans, and *r* is the human recovery rate. For each temperature-sensitive trait in each mosquito species, we fit either symmetric (Quadratic, -*c*(*T* − *T*_0_)(*T* − *T_m_*)) or asymmetric (Brière, *cT*(*T* − *T*_0_)(*T_m_* – *T*)^1/2^) unimodal thermal response models to the available empirical data [47]. In both functions, *T*_0_ and *T_m_* are respectively the minimum and maximum temperature for transmission, and *c* is a positive rate constant.

We consider a normalized version of the *R*_0_ equation such that it is rescaled to range from zero to one with the value of one occurring at the unimodal peak. Although absolute values of *R*_0_ that are used to determine when transmission is stable depend on additional factors not captured in our model, the minimum and maximum temperatures for which *R_0_* > 0 map exactly onto our normalized equations, allowing us to accurately calculate whether or not transmission should be possible at all. Empirical estimates of absolute values of *R_0_* are difficult to obtain in any case, but it is much easier to determine whether transmission is occurring and for how long. While different model formulations for predicting *R_0_* versus temperature can produce results with different magnitudes and potentially different overall shapes [48], the temperatures for which *R_0_* is above or below zero (or one) are mostly model independent. For instance, two competing models differ only by whether or not the formula in equation (1) is squared, but the square of a number (e.g., an absolute *R_0_* value) greater than one is always greater than one, and the square of a number less than one is always less than one. Therefore, the threshold temperatures at which absolute *R_0_* > 0 or absolute *R_0_* > 1 will be exactly the same for either choice of formula (Fig. S6). Similarly, because different expressions for *R_0_*, including the square of equation (1), map monotonically onto our function, they will produce identical estimates for the temperatures at which transmission declines to zero and peaks (Fig. S6). Consequently, our use of relative *R_0_* adequately describes the nonlinear relationship between mosquito and virus traits and transmission.

We fit the trait thermal responses in equation (1) based on an exhaustive search of published laboratory studies that fulfilled the criterion of measuring a trait at three or more constant temperatures, ideally capturing both the rise and the fall of each unimodal curve (Tables S1-S2). Constant-temperature laboratory conditions are required to isolate the direct effect of temperature from confounding factors in the field and to provide a baseline for estimating the effects of temperature variation through rate summation [49]. We attempted to obtain raw data from each study, but if they were not available we collected data by hand from tables or digitized data from figures using WebPlotDigitizer [50]. We obtained raw data from Delatte [19] and Alto [21] for the *Ae. albopictus* egg-to-adult survival probability (*pEA*), mosquito development rate (MDR), gonotrophic cycle duration (GCD, which we assumed was equal to the inverse of the biting rate) and total fecundity (TFD) (Table S2). Data did not meet the inclusion criterion for CHIKV or ZIKV vector competence (*b, c*) or extrinsic incubation period (*EIP*) in either *Ae. albopictus* or *Ae. aegypti*. Instead, we used DENV EIP and vector competence data, combined with sensitivity analyses.

Following Johnson *et al*. [51], we fit a thermal response for each trait using Bayesian models. We first fit Bayesian models for each trait thermal response using uninformative priors (*T_0_* ~ Uniform (0, 24), *T_m_* ~ Uniform (25, 45), *c* ~ Gamma (1, 10) for Brière and *c* ~ Gamma (1, 1) for Quadratic fits) chosen to restrict each parameter to its biologically realistic range (i.e., *T_0_* < *T_m_* and we assumed that temperatures below 0°C and above 45°C were lethal). Any negative values for all thermal response functions were truncated at zero, and thermal responses for probabilities (*p_EA_, b*, and *c*) were also truncated at one. We modeled the observed data as arising from a normal distribution with the mean predicted by the thermal response function calculated at the observed temperature, and the precision *τ*, (*τ* = 1/*σ*), distributed as *τ* ~ Gamma (0.0001, 00001). We fit the models using Markov Chain Monte Carlo (MCMC) sampling in JAGS, using the R [52] package *rjags* [53]. For each thermal response, we ran five MCMC chains with a 5000-step burn-in and saved the subsequent 5000 steps. We thinned the posterior samples by saving every fifth sample and used the samples to calculate *R_0_* from 15-40°C, producing a posterior distribution of *R_0_* versus temperature. We summarized the relationship between temperature and each trait or overall *R_0_* by calculating the mean and 95% highest posterior density interval (HPD interval; a type of credible interval that includes the smallest continuous range containing 95% of the probability, as implemented in the *coda* package [54]) for each curve across temperatures.

We fit a second set of models for each mosquito species that used informative priors to reduce uncertainty in *R_0_* versus temperature and in the trait thermal responses. In these models, we used Gamma-distributed priors for each parameter *T_0_, T_m_, c*, and *τ* fit from an additional ‘prior’ dataset of *Aedes* spp. trait data that did not meet the inclusion criteria for the primary dataset (Table S3). We found that these initial informative priors could have an overly strong influence on the posteriors, in some cases drawing the posterior distributions well away from the primary dataset, which was better controlled and met the inclusion criteria. We accounted for our lower confidence in this data set by increasing the variance in the informative priors, by multiplying all hyperparameters (i.e., the parameters of the Gamma distributions of priors for *T_0_, T_m_*, and *c*) by a constant *k* to produce a distribution with the same mean but 1/*k* times larger variance. We chose the value of *k* based on our relative confidence in the prior versus main data. Thus we chose *k* = 0.5 for *b, c*, and *PDR* and *k* = 0.01 for *lf*. This is the main model presented in the text (Fig. 2). It is comparable to some but not all previous mechanistic models for *Ae. aegypti* and *Ae. albopictus* transmission (Fig. S5). Results of our main model, fit with informative priors, did not vary substantially from the model fit with uninformative priors (Figs. S7-S8).

### Incorporating daily temperature variation in transmission models

Because organisms do not typically experience constant temperature environments in nature, we incorporated the effects of temperature variation on transmission by calculating a daily average *R_0_* assuming a daily temperature range of 8°C, across a range of mean temperatures. This range is consistent with daily temperature variation in tropical and subtropical environments but lower than in most temperate environments. At each mean temperature, we used a Parton-Logan model to generate hourly temperatures and calculate each temperature-sensitive trait on an hourly basis [55]. We assumed an irreversible high-temperature threshold above which mosquitoes die and transmission is impossible [56,57]. We set this threshold based on hourly temperatures exceeding the critical thermal maximum (*T_m_* in Tables S1-S2) for egg-to-adult survival or adult longevity by any amount for five hours or by 3°C for one hour. We averaged each trait over 24 hours to obtain a daily average trait value, which we used to calculate relative *R_0_* across a range of mean temperatures. We used this model in the validation against human cases (Fig. 3) and the risk map (Fig. 4).

### Model validation with DENV, CHIKV, and ZIKV incidence data

To validate the model, we used data on human cases of DENV, CHIKV, and ZIKV at the country scale and mean temperature during the transmission window. Using statistical models (as described below), we estimated the effects of predicted *R_0_*(*T*) on the probability of local transmission and the magnitude of incidence, controlling for population size and several socioeconomic factors. We downloaded and manually entered Pan American Health Organization (PAHO) weekly case reports for DENV and CHIKV for all countries in the Americas (North, Central, and South America and the Caribbean Islands) from week 1 of 2014 to week 8 of 2015 for CHIKV and from week 52 of 2013 to week 47 of 2015 for DENV (www.paho.org). ZIKV weekly case reports for reporting districts (e.g., provinces) within Colombia, Mexico, El Salvador, and the US Virgin Islands were available from the CDC Epidemic Prediction Initiative (https://github.com/cdcepi/) from November 28, 2015 to April 2, 2016. We aggregated the ZIKV data into country-level weekly case reports to match the spatial resolution of the DENV, CHIKV, and covariate data.

### Temperature data collection

We matched the DENV, CHIKV, and ZIKV incidence data with temperature using daily temperature data from METAR stations in each country, averaged at the country level by epidemic week. A previous study found a six-week lagged relationship between temperature and oviposition for *Aedes aegypti* in Ecuador [39]. Assuming that the subsequent transmission, disease development, medical care-seeking, and case reporting in humans takes an additional four weeks, we assumed *a priori* a ten-week lag between temperature and incidence (i.e., mean temperature for the week that is ten weeks prior to each case report). METAR stations are internationally standardized weather reporting stations that report hourly temperature and precipitation measures. Outlier weather stations were excluded if they reported a daily maximum temperature below 5°C or a daily minimum temperature above 40°C during the study period, extremes that would certainly eliminate the potential for transmission in a local area. Because case data are reported at the country level, we needed a collection of weather stations in each country that accurately represent weather conditions in the areas where transmission occurs, excluding extreme areas where transmission is unlikely. For the study period of October 1, 2013 through April 30, 2016, we downloaded daily temperature data for each station from Weather Underground using the *weatherData* package in R [58]. We removed all data from Chile because it spans so much latitude and the terrain is so diverse that its country-level mean is unlikely to be very representative of the temperature where an outbreak occurred.

### Socioeconomic covariate data

We accessed available data on projected 2016 gross domestic product (GDP) for countries of interest via the International Monetary Fund’s World Economic Outlook Database (http://www.imf.org/external/ns/cs.aspx?id=28). The direct and total contributions of tourism to GDP in 2016 were compiled from World Travel and Tourism Council economic impact reports (http://www.wttc.org/research/economic-research/economic-impact-analysis/country-reports/#undefined). We retrieved population size data for 2013-2015 from the United Nations Population Division (https://esa.un.org/unpd/wpp/Download/Standard/Population/) and averaged them across the three years for each country. Throughout the analyses below, unless otherwise specified, we used the natural log of the population size and of GDP as our predictors. We have two reasons for this choice. The first is that, intuitively, the relative order of magnitude of the population/GDP is more important in determining observed outbreak sizes or probabilities than their absolute sizes. Second, population sizes and GDPs across countries tend to exhibit clumped patterns with a few outliers that are much larger than the others. From a statistical perspective, using the un-transformed populations (or GDPs) results in those few large/rich countries having very high leverage in the analysis, and thus potentially skewing the results. Taking a log of the population better balances these predictors and is the standard accepted approach when using these kinds of predictors in regression models.

### Validation analyses with human incidence versus temperature datasets

To validate the *R_0_*(*T*) model while controlling for population and socio-economic factors, we used generalized linear regression on the weekly case count data. Importantly, we focused on testing whether the case counts were consistent with the transmission – temperature relationship predicted from our model, rather than on maximizing the variation explained in the statistical model. We are more specifically interested in understanding autochthonous transmission (i.e., locally acquired, not just imported cases). We set country-level thresholds for the number of cases defining autochthonous transmission for our three diseases separately, based on current transmission understanding: seven cases of CHIKV, 70 cases of DENV, and three cases of ZIKV. We derived these thresholds in the following way. First, we looked for data on outbreaks of travel related cases in countries that are not expected to experience any local transmission. For instance, in 2014 Canada experienced 320 confirmed, travel-related cases of chikungunya (http://www.phac-aspc.gc.ca/publicat/ccdr-rmtc/15vol41/dr-rm41-01/rapid-eng.php), equivalent to an average of more than six cases per week. Thus, to be conservative in our estimates, we set the threshold of transmission as seven cases/week for CHIKV. The reported weekly cases of DENV transmission in our study sample are considerably higher than for CHIKV (mean DENV incidence was nearly 100 times higher mean CHIKV incidence). We chose a moderately high threshold of 70 cases in a week (i.e., 10 times higher than the CHIKV threshold based on Canadian cases) to reflect higher overall incidence and increased potential for travel related cases. We examined the sensitivity of the results to choice of threshold by varying it from 25 to 100, and we found qualitatively similar results for all thresholds that we tested. As ZIKV is not as well established as either CHIKV or DENV at this time, smaller numbers of cases may indicate autochthonous transmission. Consequently, we chose a threshold of three cases for ZIKV (approximately half the CHIKV threshold). Further, the results were fairly sensitive to the ZIKV threshold as many locations have small numbers of cases. Since higher thresholds exclude a very large proportion of available case data making analysis impossible, we used the slightly less conservative threshold of three cases for autochthonous transmission of ZIKV. The resulting data consisted of zeros for no transmission and positive case counts when transmission is presumed to be occurring. To model these data, we used a hurdle model that first uses logistic regression on the presence/absence of local transmission data to understand the factors correlated with local transmission occurring or not (PA analysis). Then we modeled the log of incidence (number of new cases per reporting week) for positive values with a gamma generalized linear models (GLM; i.e., incidence analysis).

We were interested in understanding whether *R_0_*(*T*) was an important predictor of human transmission occurrence and incidence, after controlling for potentially confounding factors like population size and socioeconomic conditions. To do this, we fit a series of models with different subsets of predictors that included *R_0_*(*T*), the socioeconomic variables with population, or both (see Table S4 for full models). To control for human population size, we created new metrics based on *R_0_*(*T*) and population size to use for validation against the PAHO incidence data. We define *GR_0_*, which is the posterior probability that *R_0_*(*T*) > 0. We use *log*(*p*)*\*GR_0_*, where *p* is the population size, as the relevant *R_0_*-based predictor for the PA analysis. For the incidence analysis, we instead use *log*(*p*R_0_*(*T*)) as the predictor. In all cases log refers to the natural logarithm. For simplicity, we refer to these as the *R_0_*(*T*) metrics hereafter and in the Results.

In both the PA and incidence analyses, we first used the full data sets to examine which of the candidate models best described the data. Randomized quantile residuals indicated that the logistic and gamma GLM models were performing adequately. We compared the approximate model probabilities, calculated from the BIC scores, as well as the proportion of deviance explained (D^2^) from each model. Next we examined the performance of the models in predicting out of sample, for both PA and incidence analyses. To do this we created 1000 random partitions, where 90% of the data were used to train the model and 10% were used for testing. In the PA analyses we classified each partition based on presence/absence, with separate classification thresholds for DENV versus CHIKV/ZIKV as these grouping had much different probabilities of occurrence. We assessed the performance of the model for the PA analysis based on the mean misclassification rate. In the incidence analyses we assessed the model performance based on the predictive mean absolute percentage error (MAPE). Since differences in prediction success between the models in both the PA and incidence analyses were not statistically significant, we present the simpler models that only include the *R_0_*(*T*) metrics in the main text (Fig. 3) and the models that additionally include socioeconomic covariates in the Supplementary Information (Figs. S2-S3). We plotted the model predictions as a function of the *R_0_*(*T*) metrics together with the observed data for the PA and incidence analyses using the R package *visreg* [59].

The residuals of the incidence model exhibit “inverse trumpeting,” in which residual variation is larger at low than high predicted incidence (Fig. S9). This occurs in part because we forced the model to go through the origin, i.e., no transmission when *R_0_*(*T*) or the population size is equal to zero. However, the data did sometimes show transmission where we did not expect it, potentially because of imported cases, errors in reporting, or small pockets of transmission suitability in countries or times that are otherwise unsuitable on average. More local-scale case reporting that separates autochthonous from travel-associated cases would be needed to tease apart the source of this error.

### Mapping temperature suitability for transmission

Using our validated model, we were interested in where the temperature was suitable for *Ae. aegypti* and/or *Ae. albopictus* transmission for some or all of the year to predict the potential geographic range of outbreaks in the Americas. We visualized the minimum, median, and maximum extent of transmission based on probability of occurrence thresholds from the *R_0_* models for both mosquitoes. We calculated the number of consecutive months in which the posterior probability of *R_0_* > 0 exceeds a threshold of 0.025, 0.5, or 0.975 for both mosquito species, representing the maximum, median, and minimum likely ranges, respectively. The minimum range is shown in Fig. 4 and all three ranges are overlaid in Fig. S4. This analysis indicates the predicted seasonality of temperature suitability for transmission geographically, but does not indicate its magnitude. To generate the maps, we cropped monthly mean temperature rasters from 1950-2000 for all twelve months (Worldclim; www.worldclim.org/) to the Americas (*R, raster* package, *crop* function) and assigned cells values of one or zero depending on whether the probability that *R_0_* > 0 exceeded the threshold at the temperatures in those cells. We then synthesized the monthly grids into a single raster that reflected the maximum number of consecutive months where cell values equaled one. The resulting rasters were plotted in ArcGIS 10.3, overlaying the three cutoffs (Fig. S4). We repeated this process for both mosquito species.

## Acknowledgments

Barry Alto, Krijn Paaijmans, Francis Ezeakacha, and Helene Delatte kindly provided raw data used in the analyses. We gratefully acknowledge the Centers for Disease Control and Prevention Epidemic Predictions Initiative (CDC EPI) for collating and sharing the Zika incidence data on GitHub (https://zenodo.org/record/48946#.Vz-EM2bb8ys).

## Supporting Information Legends

**S1 Appendix. Supplementary Results, References, Figures S1-S15, and Tables S1-S3.**

**S2 Appendix. Table S4.** Model validation results.

**S3 Appendix. Online code, data, and analyses.** Available as a ZIP file on Figshare: https://figshare.com/s/b79bc7537201e7b5603f, DOI: https://dx.doi.org/10.6084/m9.figshare.4563928.

